# Activity-Dependent Remodeling of Corticostriatal Axonal Boutons During Motor Learning

**DOI:** 10.1101/2024.06.10.598366

**Authors:** Mengjun Sheng, Di Lu, Kaiwen Sheng, Jun B Ding

## Abstract

Motor skill learning induces long-lasting synaptic plasticity at not only the inputs, such as dendritic spines^1-4^, but also at the outputs to the striatum of motor cortical neurons^5,6^. However, very little is known about the activity and structural plasticity of corticostriatal axons during learning in the adult brain. Here, we used longitudinal in vivo two-photon imaging to monitor the activity and structure of thousands of corticostriatal axonal boutons in the dorsolateral striatum in awake mice. We found that learning a new motor skill induces dynamic regulation of axonal boutons. The activities of motor corticostriatal axonal boutons exhibited selectivity for rewarded movements (RM) and un-rewarded movements (UM). Strikingly, boutons on the same axonal branches showed diverse responses during behavior. Motor learning significantly increased the fraction of RM boutons and reduced the heterogeneity of bouton activities. Moreover, motor learning-induced profound structural dynamism in boutons. By combining structural and functional imaging, we identified that newly formed axonal boutons are more likely to exhibit selectivity for RM and are stabilized during motor learning, while UM boutons are selectively eliminated. Our results highlight a novel form of plasticity at corticostriatal axons induced by motor learning, indicating that motor corticostriatal axonal boutons undergo dynamic reorganization that facilitates the acquisition and execution of motor skills.

## Main

Learning and executing fine movement skills require corticostriatal circuits^7-10^. During motor learning, neuronal ensembles in the primary motor cortex (M1) first expand the movement-related population and then refine it into a smaller population that generates reproducible spatiotemporal sequences of activity^8,11^. Motor cortical neurons project to the dorsolateral striatum (DLS)^12-14^ and drive the activity of striatal spiny projection neurons (SPNs)^15^. Neuronal activity in the DLS also reflects those of motor cortical neurons during motor learning^15^ and reorganizes into a stable pattern that encodes the sequences of motion^16^. Mechanistically, motor learning leads to the selective remodeling of dendritic spines, where new glutamatergic synapses are formed, and the strengthening of their outputs to the striatum^6^. Conversely, in movement disorders such as Parkinson’s disease, the SPN dendritic spines are diminished, impeding corticostriatal synaptic transmission^17-19^. Similar to postsynaptic dendritic spines, their presynaptic partners – axonal boutons – signal synapse formation and elimination through their addition and subtraction^20-23^. However, it is unknown whether and how corticostriatal axons and boutons undergo activity and morphological alternations *in vivo* in the adult brain following motor learning.

We trained mice to perform a cued lever-pushing task under a two-photon microscope (Fig.1a), as described previously^16^. Briefly, a lever push beyond a set threshold after the cue onset was rewarded with water. An inter-trial interval (ITI) was introduced between consecutive trials, during which any un-cued lever push was not rewarded and triggered an additional timeout (see Methods). Mice were trained daily with this task for approximately 2 weeks (n=17 mice). The success rate of achieving a reward (Fig.1b) and the timing of their reaction (Fig.1c) improved throughout the training. After an initial increase in lever pushes during ITI, the mice learned to curb the unrewarded pushes and decreased the overall number of lever-pushing during ITI in later sessions (Fig.1d). Importantly, lever movement trajectories on individual trials became more stereotyped over time (Fig.1e) with higher pair-wise correlation across later sessions (Fig.1f), a hallmark of learning a new motor skill^8,11,16^.

**Figure 1:**
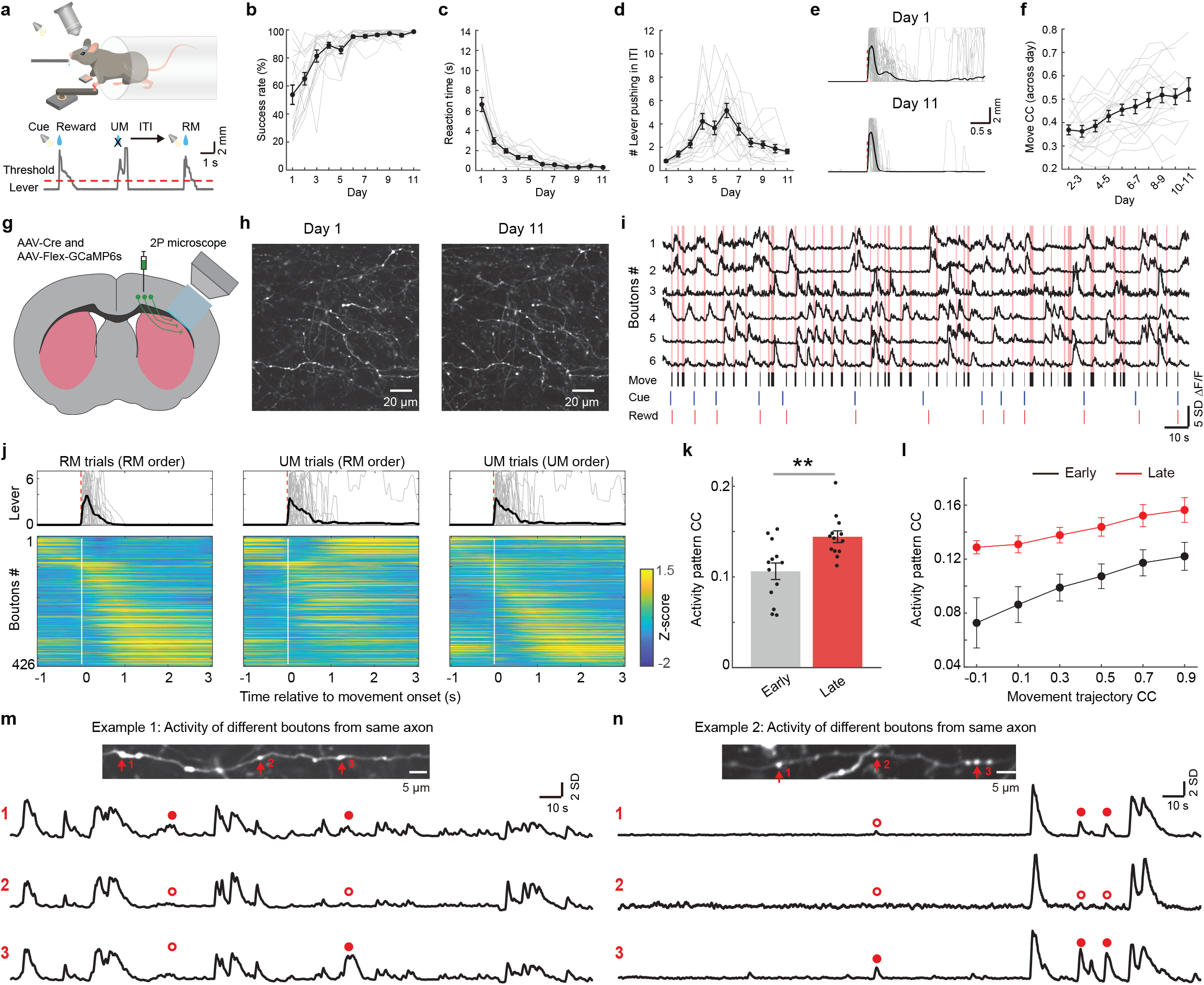
Longitudinal two-photon Ca^2+^ imaging of corticostriatal axonal boutons during motor learning. **a**, Task schematic. Mice were trained to push the lever to obtain water reward after the cue onset. This example shows two rewarded pushes (RM) and one un-rewarded push (UM) during ITI. **b**, The increase in success rate, and **c**, the decrease in reaction time, and **d**, the number of lever pushing during ITI (inter-trial-interval, n=17 mice). Grey, the performance of individual mice, black, average of all mice. **e**, Representative lever pushing movement trajectories in rewarded trials from one mouse at the early stage (day 1) and late stage (day 11), Grey, individual trials, black, average of all trials; red dotted line, movement onset. **f**, pairwise movement correlation on individual trials across sessions. Movement trajectories became more similar across trials (r=0.44, P=1.05*10^−9^, Pearson’s correlation). **g**, Schematic diagram showing the sites of virus injection (M1) and imaging (dorsolateral striatum, DLS). **h**, Representative GCaMP6s images of corticostriatal axons in DLS imaged on day 1 and day 11. **i**, Examples of task-related activities of corticostriatal axonal boutons from one mouse. Vertical red line: lever movement period; black, lever movement period; blue, cue; red, reward. (Horizontal bar, 10 s; vertical bar, 5 SD Z-scores ΔF/F). **j**, Top: individual (grey) and average (black) lever pushing trajectories in rewarded movements (RM) and unrewarded movements (UM). Bottom, corresponding averaged activity of 426 axonal boutons. Activity was aligned by either RM (left), or UM (middle and right) onset time (white line). Boutons were sorted according to the order of the time of averaged peak activity in RM (left and middle), or UM (right) trials. **k**, Pair-wise correlation on trial-to-trial activity during RM trials in early and late stages of motor learning (P= 0.007, two-sided Wilcoxon rank sum test, n=13 mice). **l**, Pair-wise trial-to-trial correlation plotted as a function of movement correlation on those trials. A more robust relationship between activity and movements appeared during learning (P=0.001,0.007,0.006,0.004,0.014 and 0.046 for bins 1 to 6 between early and late stage, respectively, two-sided Wilcoxon rank sum test, n=13 mice). **m-n**, Top: Examples of averaged GCaMP6s images showing clear bouton structures localized on the same axon. Bottom: Examples ΔF/F_0_ traces from 3 different boutons (red arrow) localized on the same axon. Red dots: bouton-specific local Ca^2+^ events. Horizontal bar, 10 s; vertical bar, 2 SD Z-scores Δ F/F_0_. Error bars represent SEM.

### Movement-related M1 axonal bouton activities

To investigate how the activities of the motor cortical axonal boutons are modified during motor learning, we combined the lever-pushing task with longitudinal *in vivo* two-photon calcium imaging. Before training, we injected adeno-associated virus (AAV) encoding the genetically encoded Ca^2+^ indicator, GCaMP6s^24^, into layer 5 of the forelimb region of the M1^25^ and implanted a chronic imaging window above the DLS (Fig.1g, see Methods). Approximately 2 to 4 weeks after surgery, we imaged M1 axons and boutons in the DLS through the chronic window while simultaneously monitoring the mouse’s behavior.

We repeatedly imaged the activities of populations of M1 axons and boutons as mice learned and performed the lever-pushing task (Fig.1h). High correlations of bouton activity and lever-push movements were evident at the single bouton level (Fig.1i). Previous studies have shown that M1 movement-related neurons displayed reproducible activity timing relative to movement onset in later sessions^11^. Analyzing if axonal activity followed a similar pattern, we found that the activity sequence spanned over the entire duration of the rewarded movement (RM, Fig.1j left), consistent with M1 somatic activity. Interestingly, despite the movement trajectories looking similar to those of the RM’s, the temporal activity pattern of the same ensemble of boutons during un-rewarded movement (UM) trials looked markedly different (UM activity ordered by peak activity timing in RM trials, Fig.1j middle). However, if the boutons were re-sorted according to the order of the time of averaged peak activity in UM trials, a similar temporal activity pattern emerged again (Fig.1j right). Consistent with previous reports^11^, the pair-wise correlation on trial-to-trial activity significantly increased in the late stages of motor learning compared to that of the early stages (Fig.1k, early stage: day 1 to day 3; late stage: >= day 8). We further evaluated the relationship between movement and axonal bouton activity during early and late stages of learning. By sorting trials according to the similarity of movements for each pair of trials, we found the overall activity pattern pair-wise correlation was significantly higher in the late stage compared to the early stage, even when mice generated dissimilar movement trajectories (Fig.1l), suggesting that the overall bouton activity pattern became more reproducible at late sessions regardless of movement similarity.

Cortical efferent axons arbor extensive collaterals in the striatum and form *en passant* synapses with postsynaptic striatal neurons^26,27^. This is evident in our data set – a subset of boutons that belong to the same parent axon could be clearly identified (Fig.1m-n). Because of the high fidelity of action potential propagation along the axons^28-30^, multiple release sites at these *en passant* synapses are thought to deliver cortical outputs faithfully to multiple postsynaptic striatal neurons. Upon careful examination of the activity of the subsets of boutons on the same stretch of an axon, unexpectedly, we observed heterogeneous responses – while most Ca^2+^ transients were uniformly present in all boutons observed, there were localized additions or failures of activities at discrete axonal boutons (Fig.1m-n).

Together, our results suggest that the activity of M1 corticostriatal axonal boutons is movement-related and can be modulated by reward. Furthermore, the boutons formed on the same axons show heterogeneous activity patterns.

### Reward modulation of movement-related bouton activities

To further characterize the reward modulation of bouton activities, we asked whether the individual movement-related bouton activities differ based on their responses to reward outcomes. When we aligned the time-varying GCaMP6s signal of all trials to the onset of either RM or UM trials, we saw a clear distinction. Specific boutons had reproducible temporal activity patterns in RM trials but not in UM trials (Fig.2a left); conversely, some boutons were only active during UM trials but not RM trials (Fig.2a right), while some boutons were active during lever pushing movement regardless of reward outcome (Fig.2a middle). Therefore, we classified the boutons into three categories: RM-only, UM-only, and RM-UM both boutons (see Methods). To simplify, in some analyses, RM-only and RM-UM boutons were grouped as RM-responsive boutons. To further confirm our classification unbiasedly, we performed a Principal Component Analysis (PCA) on RM and UM trials (see Methods) and asked whether RM-only and UM-only bouton activity could be discernible in a low-dimensional space. Briefly, we averaged the RM and UM trials for each bouton and then concatenated, creating a 2M-by-N matrix, where M denoted the number of time points per RM or UM trial (ranging from –1 to 3 s relative to movement onset), and N represented the number of boutons. We then extracted the first three principal components, visualizing the RM and UM trials in a 3-dimensional principal component (PC) space. Each bouton occupied a unique position as a distinct dot in this space (Fig.2b). The RM-only (red dots) and UM-only (blue dots) boutons exhibited clear separation in the 3-D PC space.

**Figure 2:**
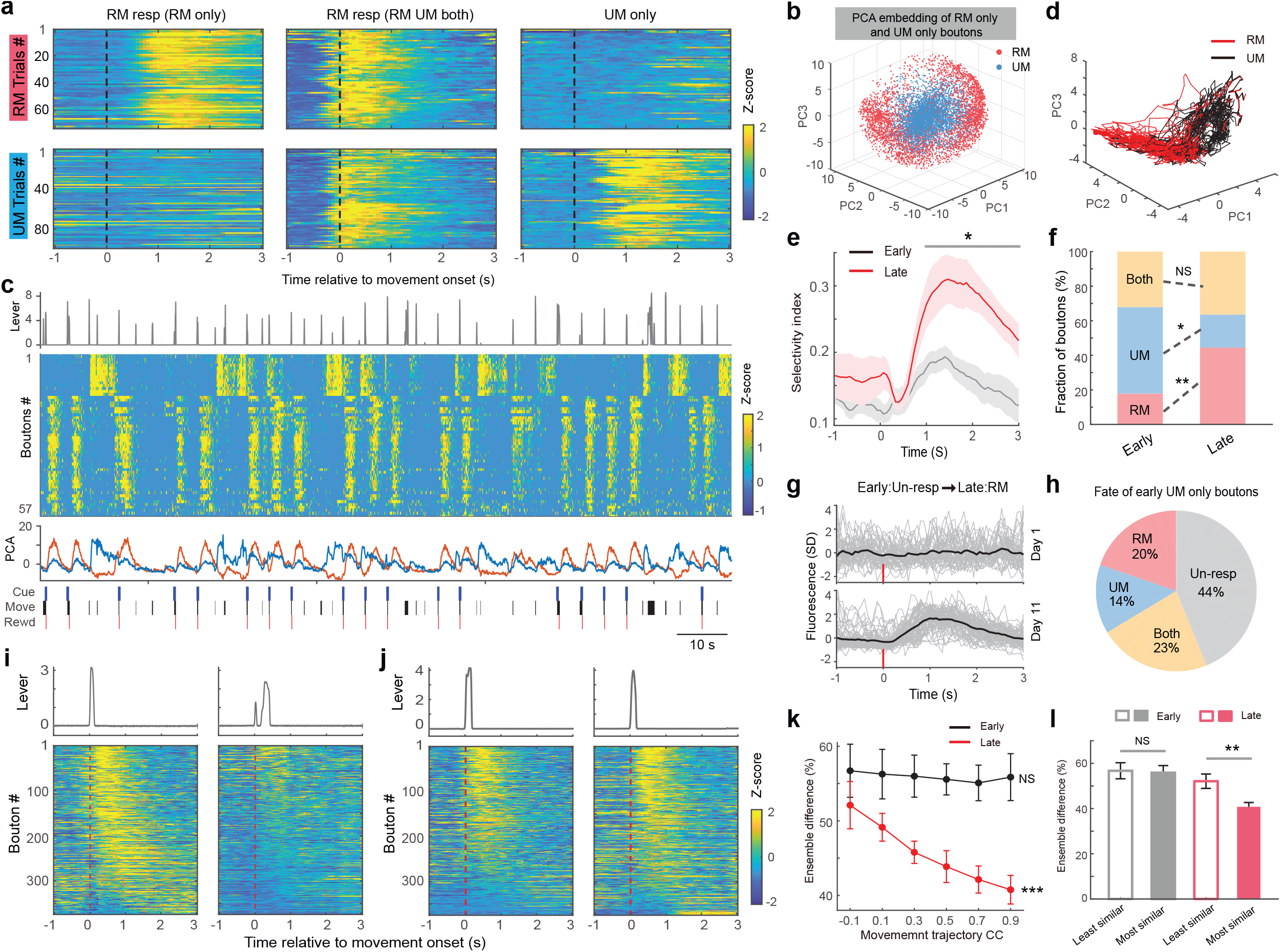
Reward and movement execution related M1 boutons. **a**, Examples of the activity of 3 individual boutons during RM (top) and UM (bottom) trials. The peri-movement activity of the boutons shows reward selectivity. Left: bouton only responded in RM trials; middle, bouton was active during both RM and UM trials; right, bouton was only activated during UM trials. **b**, PCA embedding of all corticostriatal boutons (n=3744 RM boutons and n= 4211 UM boutons form 8 mice). Red, RM only boutons; blue, UM only boutons. **c**, Top: Lever movement trajectory (unit: mm). Second, Average activity of simultaneously imaged UM (1-15) and RM (16-57) boutons from one mouse. Each row represents a bouton. Third, PC1 (orange) and PC2 (blue) for population bouton activity. Bottom, annotations of behavior: blue, cue; black, lever movement; red, reward. **d**, Three-dimensional (first three PCs) neural activity trajectories of RM (red) and UM (black) trials from one representative session. **e**, Trajectory selectivity index for RM or UM trials at early (black) and late stage (red) of learning (P<0.05, two-sided Wilcoxon rank sum test, n=8 mice, see methods for details). The shaded area represents SEM. **f**, Change of bouton reward selectivity during motor learning. RM, reward movement responsive boutons; UM unrewarded movement selective boutons; both, boutons responsive to both RM and UM. (*P<0.05, **P<0.01, NS, not significant, two-sided Wilcoxon rank sum test, n= 8 mice). **g**, Example bouton became active during RM trials. **h**, Late fate of UM boutons during the early stage of motor learning. The reward selectivity of the boutons changed throughout motor learning. **i-j**, Examples of trials showing movement trajectories (top) and activity of bouton ensembles (bottom) in dissimilar (**i**) and similar (**j**) activated RM ensembles. **k**, Difference in the percentage of activated bouton ensemble plotted against the pair-wise movement correlation. Higher movement correlation pairs show smaller differences in activated bouton ensembles, while dissimilar movement pairs show larger differences in activated bouton ensembles at the late stage of learning (Early: r=-0.04, P=0.73; Late: r=-0.46, P=1.89*10^−5^, Pearson’s correlation, n=13 mice). Error bars represent SEM. l, Ensembles difference between trials with most similar or least similar movement trajectories (**P<0.01, NS, not significant, two-sided Wilcoxon rank sum test, n= 13 mice).

We next investigated the relationship between PCs, bouton activity, and behavior outcome. To do so, we examined the population bouton activity of RM-only and UM-only boutons in consecutive trials, plotted the amplitude of PC over time, and aligned with movement behavior (Fig.2c). We found that PC1 could faithfully depict the RM-related responses while PC2 represented the UM-related responses. In addition, we plotted the activity trajectories of individual RM and UM trials and found that RM and UM activity trajectories were largely separated in the 3D low dimensional space (Fig.2d). Lastly, to measure the separation of RM and UM activity trajectories in the 3D PCA space, we calculated the selectivity index for RM and UM trials (see Methods). We found that, compared to the early stage, the selectivity index was significantly increased at the late stage of motor learning, indicating that the bouton population activity became more selective to reward (Fig.2e).

The emergence of reproducible timing activity patterns of corticostriatal boutons in late sessions may result from reward-based reinforcement of certain activity-reward outcome pairs out of initial exploration during learning. In this case, the activity of RM-only or UM-only boutons during the early stage may have a similar representation at the late stage. Alternatively, the learned activity pattern may require dynamic rearrangement of bouton ensembles, which may result in changes in the representation of RM or UM. To distinguish these possibilities, we calculated the proportions of RM-only, UM-only, and RM-UM both boutons, at the early and late sessions. We found that the fraction of UM-only boutons was significantly decreased, the fraction of RM-only boutons was significantly increased, and the percentage of RM-UM both boutons remained the same (Fig. 2f). To further reveal the dynamic change of bouton representation of RM and UM, we meticulously tracked the activity of the same boutons during the early and late stages (Fig.2g) and analyzed the fate of classified boutons at early stage, and the origin of the classified boutons at late stage (Fig.2h). We found that only ∼35% of the boutons maintained their stable representation, and the majority of the boutons changed their representation of reward outcome. For example, nearly half of the UM-only boutons at early sessions became unresponsive at late sessions, and ∼20% of them became RM-only boutons (Fig. 2h).

Previous studies revealed that the M1 cortical neuron activity pattern was reproducible with learned movement only in the expert mice, whereas similar movements made in early sessions were accompanied by different activity patterns^8,11^. Because the biggest change in bouton representation after learning was the increase of RM-responsive boutons, we asked whether the activity of the RM-responsive boutons was better correlated with movement execution. We analyzed the movement trajectories and the activated RM ensembles for each trial pair (Fig.2i-j, two example trial pairs with different activated RM ensemble (i) and similar activated ensemble (j)). We found a significant relationship between the similarity of movement trajectories and the fraction of activated boutons for each trial pair (Fig.2k and I, Early stage: 1470 trials, Late stage: 2109 trials, n= 13 mice). In other words, similar fractions of activated boutons in the ensemble would result in similar and consistent movement trajectories.

Noticeably, such a relationship is only true for late sessions but not early sessions (Fig.2k and l). Together, the results indicate that motor learning stabilizes the general relationship between activity and movement in pairs of trials, which is accompanied by changes in the identity of bouton representation of reward outcome.

### Bouton-specific activity and movement behavior

Our data indicated that, at a population level, the activities of axonal boutons show heterogeneous responses to movements based on reward outcomes (RM vs UM). More surprisingly, at a single axon level, different boutons on the same axons also display heterogeneous activity patterns (Fig 1m-n). Here, we ask whether the bouton-specific activities are related to behavior outcomes and whether motor learning can further modulate activity patterns of boutons on the same axons. Therefore, we focused the analyses on the populations of boutons on the same axons and aligned their activity with behavior (Fig 3a), and consistently, most Ca^2+^ transients were related to movements (RM or UM). By aligning bouton activities between an example pair of boutons, it is clear that while most of the Ca^2+^ transients were present in both boutons, there were ample local activities that only occur in one bouton but not another (Fig.3a). To prevent bias in bouton activities influenced by the highest or the least amount of Ca^2+^ transients seen in individual boutons, we first applied detection criteria to define the Ca^2+^ peaks for each bouton. We then compared the timing of the Ca^2+^ peaks between every pair of boutons, categorizing them as either same peaks (Ca^2+^ transients detected in both boutons) or unique peaks (Ca^2+^ transients detected only in one of the boutons, see methods).

**Figure 3:**
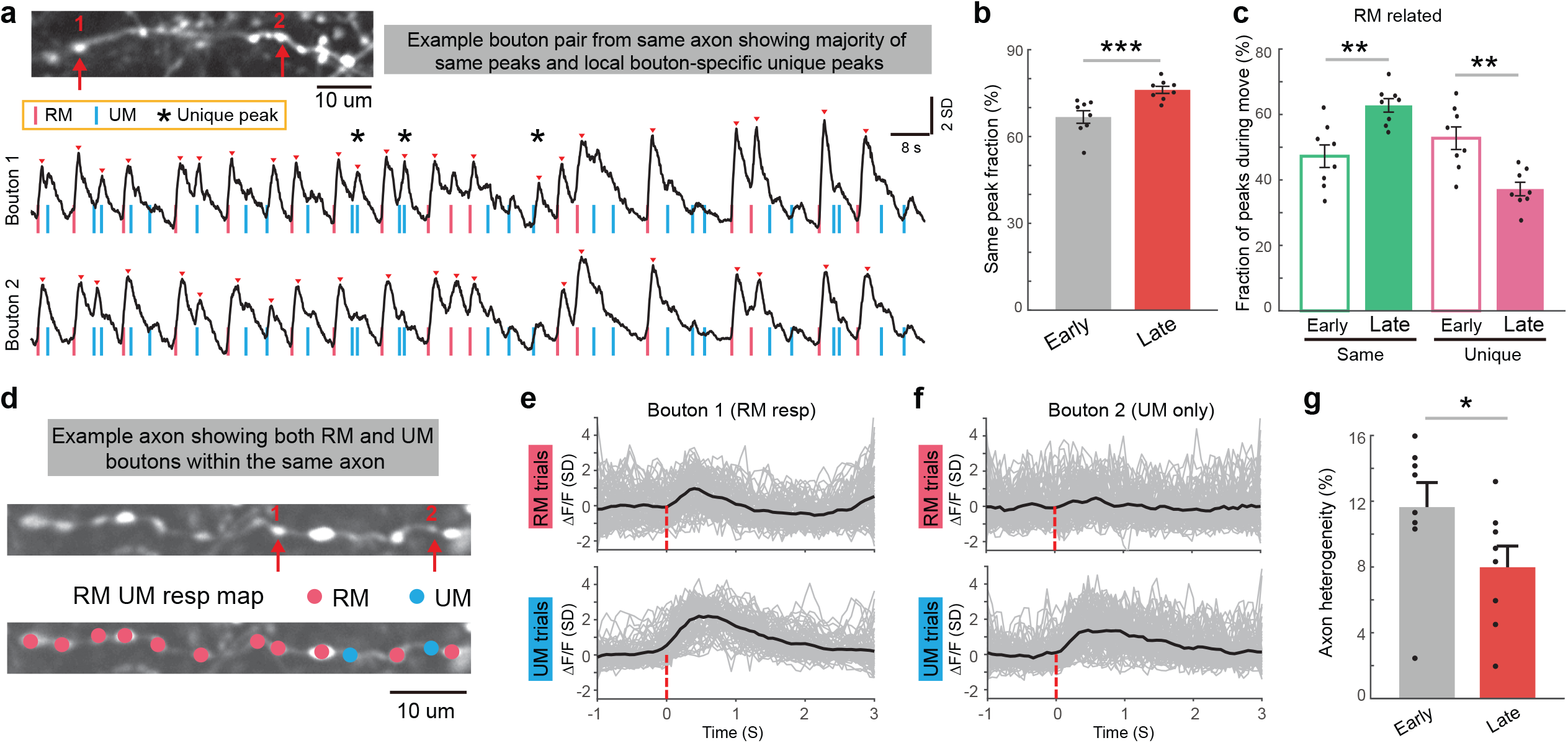
Heterogeneity in activities of boutons located on the same axon. **a**, Top: example of average GCaMP6s image showing a single axon with clear axon and bouton morphology. Bottom, representative Ca^2+^ traces of two distinct boutons (red arrow) located on the same axon. Red vertical line: initiation of RM; blue vertical line, initiation of UM; red arrowhead, detected Ca^2+^ transients; stars, heterogeneous local Ca^2+^ transients. **b**, fractions of unified Ca^2+^ transients at early and late stages of motor learning. (p=0.001, two-sided Wilcoxon rank sum test, n=8 mice). **c**, Relative fraction of RM related same peaks and unique peaks at early and late stage (**P<0.01, two-sided Wilcoxon rank sum test, n= 8 mice). **d**, Example GCaMP6s image showing clear axon and bouton structures (top), and identified boutons responsive to RM or UM trials. **e-f**, Activities of bouton 1 (**f**), and bouton 2 (**g**) shown in **d** during RM (top) and UM (bottom) trials. Grey, Ca^2+^ transients in individual trials (Δ F/F_0_); black, average of all trials in one day (day 14). Note that both RM and UM boutons exist on the same axon. **h**, Axon heterogeneity at early and late stage (*P<0.05, two-sided Wilcoxon rank sum test, n= 8 mice). Error bars represent SEM.

We first set to determine the prevalence of same peaks *in vivo* while mice performing the task. Analyzing each entire imaging segment (∼4 min) during both the early and late training periods, we observed that ∼65% Ca^2+^ transients were uniformly detected in pairs of boutons (same peaks) during the early training period. Interestingly, the percentage of the same peaks increased to ∼80% in late training sessions (Fig 3b). These data indicated that different boutons on the same axons exhibited surprisingly high heterogenous activity patterns *in vivo*, nearly ∼35% in the early phase of the training, and this heterogeneity could be reduced by motor learning. What could contribute to the change of heterogeneous activities between boutons? Because axonal bouton activity was selective to reward outcomes (Fig 2), we next focused our analyses on Ca^2+^ transients occurring during the RM trials and asked whether motor learning could modify the consistency of bouton activities formed on the same axons. When we compared the percentage of the same peaks vs unique peaks associated with RM trials in the early and late phases of training, we observed a significant increase in the fraction of the same peaks and a significant decrease in the fraction of unique peaks (Fig 3c). We found a similar result when we analyzed Ca^2+^ transients occurring during UM trials.

The presence of unique peaks among bouton pairs also raised the question of whether boutons on the same axons could be exclusively RM- or UM-responsive. To address this, we identified the Ca^2+^ transients with RM or UM and mapped their locations along the same axons (Fig 3d). Interestingly, within the same axon, boutons predominantly displayed uniform RM- or UM-selectivity. However, even an RM-dominating axon contained some UM-selective boutons (Fig 3e-f), and vice versa, a UM-dominating axon also contains RM-selective boutons. In addition, as we showed earlier (Fig 2h), at the population level, individual axonal bouton RM- or UM-selectivity could change throughout motor learning. This is also true for boutons on the same axons; the axon heterogeneity (defined by the percentage of RM- or UM-boutons throughout the axonal segment) decreased after motor learning (Fig 3g).

Together, these data demonstrated a surprisingly high occurrence of heterogeneous activities among boutons formed on the same short stretch of axons. In addition, this heterogeneity is dynamically modulated by learning throughout the training.

### Structural plasticity of corticostriatal axonal boutons during motor learning

The changes in activity patterns and reward representations of M1 corticostriatal axonal boutons indicate a dynamic regulation of corticostriatal synaptic transmission. In postsynaptic striatal spiny projection neurons, dendritic spines, where the glutamatergic corticostriatal synapses are formed^31,32^, undergo significant activity-dependent structural changes, for example, in mouse models of movement disorders^17-19^. Here, we ask whether motor learning could result in dynamic remodeling of presynaptic axonal bouton structures. To investigate the activity-dependent structural changes in axonal boutons, we used *in vivo* two-photon microscopy to repeatedly image the same corticostriatal axon segments in the DLS labeled by the expression of EGFP in the forelimb area of the M1 contralateral to the lever-pushing forelimb (see Methods). We repeatedly imaged the same axons for 11 days every other day. In the training group, the mice were trained on the cued lever-pushing task starting on day 1, and the control group underwent identical procedures, including water restriction, habituation, and water consumption from the licking port, but without training to push the lever. By comparing images taken from two time points, we identified axonal boutons as newly formed, eliminated, or stable (Fig.4a-b). We calculated total bouton numbers for each axon to assess whether bouton density changes following motor learning. We found that the bouton density was significantly increased from day 4 and persisted throughout the training period (Control: n= 146 axons, N= 9 mice, Training: n= 143 axons, N= 8 mice, Fig.4c). To further understand the process of motor learning-induced structural plasticity, we quantified the rate of newly formed and eliminated M1 axonal boutons. Motor learning induced a transient increase in the formation of boutons on day 4 (Fig.4d), accompanied by enhanced bouton elimination on day 6 (Fig.4e). Previous studies showed that newly formed dendritic spines in M1 layer 5 pyramidal neurons are preferentially stabilized during training^1,33^. To test whether newly formed M1 axonal boutons are also preferentially stabilized during training, we analyzed the fate of newly formed boutons during motor learning in control and trained mice. We found that, in trained mice, newly formed axonal boutons were significantly more stable. However, in control mice, most of the newly formed boutons were eliminated (Fig.4f). In addition, our previous work showed that motor learning led to a selective strengthening of M1 motor engram neuron outputs formed onto clustered spines of postsynaptic striatal SPN dendrites^6^. Therefore, we examined the position of newly formed and eliminated boutons along each axon and plotted the cumulative distribution of pairs of boutons formed on day 4. We found that the bouton pairs were significantly closer in space in trained mice compared to the control mice. The mean distance between newly formed boutons was significantly shorter in trained mice compared to control mice. However, the mean distance between eliminated boutons is not clustered along the axons.

**Figure 4:**
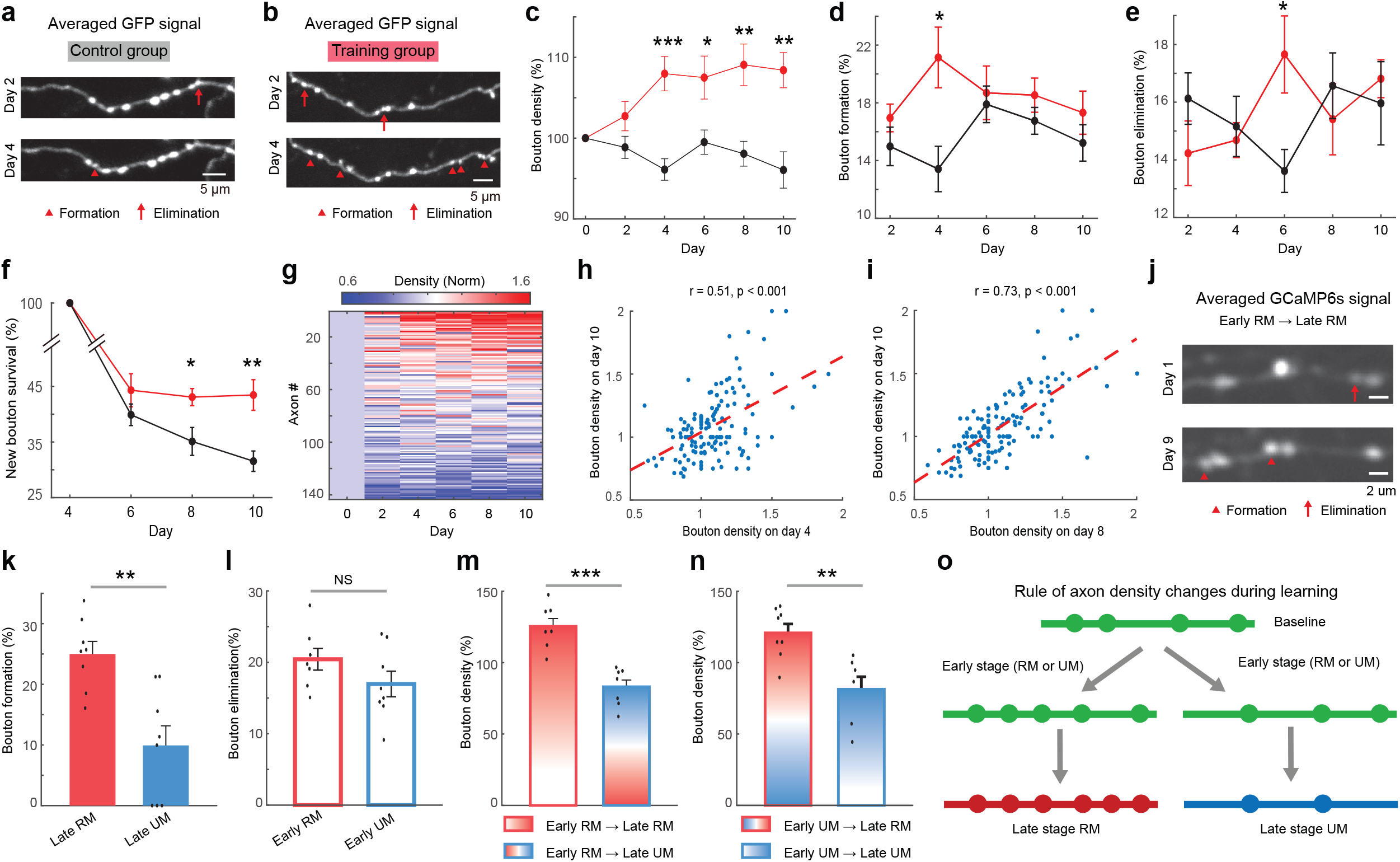
structural plasticity of corticostriatal axonal boutons during motor learning. **a-b**, Repeated imaging of the same axon (labeled with AAV-EGFP) at day 2 and day 4 reveals bouton formation (red arrowhead) and elimination (red arrow) in control (**a**) and training group (**b**). Scale bar unit: 5 µm. **c**, Changes in corticostriatal axon bouton density in the control (untrained, black) and training group mice (red). (*P<0.05, **P<0.01, ***P<0.001, two-sided Wilcoxon rank sum test, control: n=9 mice; training: n=8 mice). **d-e**, Percentage of newly formed (**d**) and eliminated (**e**) axonal boutons in control (black) and training group (red) mice. (Formation on day 4: *P<0.05, two-sided Wilcoxon rank sum test; Elimination on day 6: *P<0.05, two-sided Wilcoxon rank sum test; control: n=9 mice; trained: n=8 mice). **f**, New boutons formed on day 4 and survived, plotted as a function of time, for control (black) and training group (red) mice. (*P<0.05, **P<0.01, two-sided Wilcoxon rank sum test; control: n=9 mice; trained: n=8 mice). **g**, Changes of normalized bouton density throughout motor learning (n=143 axons from 8 mice). Axons showing higher density at the initial phase of learning tended to have higher density at the late phase of learning. **h-i**, Axonal bouton density on day 10 plotted against its density on day 4 (**h**) and day 8 (**i**). (r=0.51 and r=0.73 respectively, P<0.001, Pearson’s correlation). **j**, Images of the same axon using averaged GCaMP6s signals between early (day 1) and late (day 9) stages of learning also reveal new bouton formation (red arrowhead) and elimination (red arrow). **k**, The rate of bouton formation of those that were classified as RM (red) or UM (blue) boutons at late stage in trained mice. (**P<0.01, two-sided Wilcoxon rank sum test, n= 8 mice). **l**, The rate of bouton elimination of those that were classified as RM (red) or UM (blue) boutons at early stage in trained mice. (P=0.19, two-sided Wilcoxon rank sum test, n= 8 mice). **m**, Changes of bouton density in axons that were identified as RM axons at the early stage of training. (***P<0.001, two-sided Wilcoxon rank sum test, n= 8 mice). **n**, Changes of bouton density in axons that were identified as UM axons at the early stage (**P<0.01, two-sided Wilcoxon rank sum test, n= 8 mice). **o**, Schematic diagram showing axon density changes throughout learning. Note that regardless of whether the axons are identified as RM or UM responsive during the early stage, the axons that are responsive to RM at the late stage tend to have an increased bouton density. Conversely, axons that became UM responsive at the late stage tend to have a decreased bouton density.

The newly formed boutons form clusters along the axons, but the eliminated boutons did not have a similar spatial arrangement, indicating that bouton structural remodeling may be axon-specific. Therefore, we plotted bouton density changes throughout the learning process and sorted the axons based on their maximum density change (Fig 4g). Even though the average bouton density calculated based on all axons increased during motor learning (Fig 4c), the change in axon density diverged into two groups: axons exhibiting increased density at the early stage tended to persistently increase their density throughout the late stage, while axons decreased their density at early stage of training tended to remain lower density at the late stage (Fig 4g). When we plotted the density of each axon on day 10 against those of day 4 and day 8, it revealed a significant positive linear correlation (Fig 4h-I).

Together, these data suggest that the early stages of axonal bouton development influence final bouton density. It is possible that axons engaged in the early stages are more likely to continue to transmit corticostriatal synaptic information, and learning can further strengthen connectivity and increase synaptic transmission efficacy. Our results demonstrated that learning could change the axonal bouton selectivity to movement based on reward outcome; in particular, learning increased the proportion of RM-related boutons but reduced the UM-related ones (Fig 2f). This raised intriguing questions: are newly formed or eliminated axonal boutons activity-dependent? If so, are they dependent on the reward outcome? To address this, we used the calcium imaging data set, in which high-resolution averaged GCaMP6s images obtained from the same axons in both early- and late-stage imaging sessions could be used to clearly identify newly formed and eliminated boutons (Fig 4j) and examined their activity pattern in relation to RM or UM. Interestingly, when we calculated the bouton formation rate in RM vs UM-related axons identified at late stage, we observed a significantly higher bouton formation rate in RM axons compared to UM axons (256 RM axons, 46 UM axons, N= 8 mice, Fig 4k). However, the bouton elimination rates were similar (215 RM axons and 95 UM axons identified at early stage of learning, N= 8 mice, Fig 4l). Next, we analyzed the bouton density changes for those functionally identified axons. We found that if axons were identified as RM-related at the early stage and maintained their identity, the bouton density was higher compared to those that were identified as RM-related at the early stage and became UM-related at the late stage (Fig 4m). Conversely, axons identified as UM-related at the early stage would have a higher bouton density if they became RM-related compared to those that remained UM-selective (Fig 4n). Together, these data suggest that the formation, elimination, and maintenance of the newly formed boutons and the overall bouton density of the axons are associated with the activity of axonal boutons and dependent on behavioral outcomes (Fig 4o).

## Discussion

In the present study, we aimed to investigate how motor learning remodels corticostriatal synaptic plasticity by imaging the activity and structure of motor corticostriatal axons and axonal boutons *in vivo* in mice learning and performing a cued lever-pushing motor task. Previous studies using somatic Ca^2+^ imaging or *in vivo* recordings in the primary motor cortex revealed the formation of movement-specific cortical ensembles, whose firing covers the motion sequence and the increases in activity correlation after motor skill learning^11,34^. Our study faithfully verified these key findings now in corticostriatal axonal boutons (Fig 1i-l). In addition, we found that movement-related motor cortical axonal bouton activities in the DLS are modulated by reward outcome (Fig 2a-c). By closely examining the activities of different boutons formed on the same axon, we found surprisingly heterogeneous activity patterns on different boutons even though they are only a few microns away on the same axon (Fig 1m-n). Furthermore, these unique local heterogeneous responses are shaped by motor learning in several ways: first, motor learning can enhance the consistency of the activity responses across boutons on the same axon (Fig 3a-c); second, the bouton RM- or UM-selectivity becomes more uniform at late phases (Fig 3g); and finally, axon boutons undergo activity-dependent structural plasticity (Fig 4). Overall, corticostriatal axon and bouton activities refine throughout motor learning at both the population and single axon/bouton levels. Our longitudinal results provided a link between sub-cellular synaptic and system-level dynamics to elucidate how corticostriatal ensembles are formed and maintained throughout learning.

One of the most surprising findings is the markedly heterogeneous activity patterns among nearby boutons formed on the same axon. Decades of neuroscience research have yielded a classic model of how axons convey neuronal output to downstream postsynaptic targets^35,36^. Because of the high expression levels of voltage-gated Na^+^ and K^+^ channels^37,38^, action potential forward propagation is generally considered highly reliable and functions as a digital signal (all or none), which ensures faithful outputs^28-30,38^. In certain specialized synapses in sensory receptor cells, analog graded potential is used to increase the fidelity and capacity of synaptic information, including the rod bipolar-AII amacrine cell ribbon synapse in the retina^39^ and the hair cell ribbon synapse in the inner ear^40-42^. A combination of Analog and digital coding of axonal transmission also exists; for example, at hippocampal mossy fibers, transient subthreshold depolarizations can modulate action potential-evoked transmitter release^43^., via altering waveform of action potential (e.g. amplitude and duration)^44^. However, the heterogeneous responsive pattern revealed here represents an additional novel mechanism for information transmission at corticostriatal output. The *en passant* axonal boutons can function as a demultiplexing processer, where postsynaptic targets can receive distinct patterns of axonal output even though these targets are innervated by the same axon.

Furthermore, we demonstrated that distinct patterns of axonal bouton activity are behaviorally relevant, and motor learning can significantly increase the uniformity of bouton activity along the same axon (Fig 3). One hallmark of motor learning is the formation of stereotypic movement patterns^8,11,16,34^. In addition, motor learning reduces motion variation and jitter^8,11,16,34,45-48^. On the population level, the increased activity correlation across motor cortical neurons^11,34^, striatal neurons^16^, and here, corticostriatal axons and boutons, and the formation of stable ensembles make the corticostriatal circuits more efficient in encoding and driving movement. On the single-axon level, our revealed mechanism can also contribute to the increased efficiency, where axonal boutons became more uniform in activity patterns and RM- or UM-selectivity through activity-dependent axonal plasticity. Mechanistically, what contributes to the generation of different bouton activity patterns remains unknown. This parallel but distinct output might be due to differential inputs via axon-axonic synapses^49^. Recent studies showed that local axonal EPSPs could be evoked at dopaminergic axonal terminals in the striatum by activation of nicotinic acetylcholine receptors (nAChR)^50,51^, and conversely, activation of GABA_A_ receptor could also locally dampen axonal spikes^52^ and action potential evoked dopamine release^53^. In addition to nAChR and GABA_A_R, corticostriatal axons also express receptors of various neuromodulators, such as dopamine D1R, D2R, and endocannabinoid CB1R^54,55^. It is possible that these ionotropic and metabotropic receptors contribute to the local modulation of axonal bouton activity patterns.

## Methods

### Animals

All experimental procedures were conducted following the protocols approved by the Stanford University Animal Care and Use Committee, in accordance with the National Institutes of Health’s *Guide for the Care and Use of Laboratory Animals*. All animals were maintained on a normal 12 h:12 h light/dark cycle. WT mice (C57BL/6J, > 7 weeks) of both males and females from The Jackson Laboratory were used in the present study.

### Surgical Procedure

We performed surgeries on animals under isoflurane anesthesia (1.5% in 0.5 L/min of O_2_). To drive the expression of GCaMP6s in the motor cortex, we stereotaxically injected a mixture of AAV1-CAG-FLEX-GCaMP6s (Catalog # 100842-AAV1, 1:1) and AAV5-hSyn-Cre (Catalog # 105553-AAV5, 1:200 diluted in saline) into the caudal forelimb area of the motor cortex (from Bregma, anteroposterior (AP): 0.3 mm, mediolateral (ML): 1.5 mm; and from dura, dorsoventral (DV): -0.7 mm). Similarly, for structural imaging, we injected a mixture of AAV5-CAG-FLEX-EGFP (Catalog # 51502-AAV5, 1:1) and AAV5-hSyn-Cre (Catalog # 105553-AAV5, 1:1,000 diluted in saline). A total volume of 100-300 nL was injected over 10 minutes, using a micro pump (WPI). To prevent viral backflow, the pipette was left in situ within the brain for 15 minutes post-injection before withdrawal. Upon completion of the procedure, the incision site was sutured, and the mice were returned to their home cage once they recovered from anesthesia.

For the implantation of the chronic imaging window, 3-30 days after virus injection, we anesthetized the mice with isoflurane (1.5% in 0.5 L/min of O_2_). Following scalp removal, a titanium head plate was affixed firmly to the skull using super glue and dental cement (Lang Dental). A circular craniotomy with a diameter of approximately 2.4 mm was performed above the dorsal lateral striatum, centered at the coordinates (AP: 0.3 mm, ML: 4.0 mm). We aspirated the cortical tissue above the striatum using a 27-gauge needle at a 30-degree angle towards the surface of the corpus callosum^16,56^. Subsequently, a cannula was inserted above DLS. The cannula consisted of a stainless-steel tube (∼2.4 mm diameter, ∼1.6 mm length) and a 2.4 mm round coverslip attached to one end of the tube using adhesive (Norland optical adhesive)^16,56^. We then used Kwik-Sil and dental cement to fix the cannula and cover the exposed skull. Mice were returned to their home cage after they recovered from anesthesia.

### Two-photon imaging

*In vivo* imaging experiments were conducted using a commercial two-photon microscope (Bergamo II, Thorlabs), operated with ThorImage software. We used a 16× / 0.8 NA objective (NIKON), covering a field of view (FOV) size ranged from 120 × 120 to 200 × 200 µm (1024 × 1024 pixels). A mode-locked tunable ultrafast laser provided 925 nm excitation for two-photon imaging (Insight X3 Spectra-physics). For calcium imaging, we imaged awake mice when they were performing the lever-pushing task. Imaging data were synchronized and recorded with a PCIe-6321 card (National Instrument) to capture image frame-out timing and behavioral events, encompassing cue responses, rewards, punishments, licking behavior, and lever displacement. Time-lapse movies were acquired at an approximate frame rate of ∼15 Hz. 1 to 3 days were imaged for the early stage, 1-6 days were imaged for the late stage. For imaging the same population of axons and boutons, same FOVs were imaged between early and late stage. The first 3 days were defined as the early stage, late stage was the days when mice learned the task (>=8 days). For example, one mouse was imaged on days 1-3 and days 9-11, then day 1-3 were defined as early stage, and days 9-11 were defined as the late stage. 13 mice were used in functional calcium imaging, including 8 mice imaged the same axons and boutons at the early and late stage, another 5 mice imaged different FOVs at the early and late stage of learning.

For structural imaging, mice were anesthetized with 1% - 1.5% isoflurane and a heating pad was used to keep normothermia. Image stacks were acquired via real-time averaging of 20 frames, with a z-step of 1 μm to ensure precise axial resolution. 2-4 regions of interest (ROIs) were imaged per mouse, and these ROIs were repeatedly imaged every other day. 8 mice were used in structural imaging for the training group, and 9 mice were used for the control group.

### Cued lever-pushing task

The cued lever-pushing task was conducted as previously described^16^. Briefly, mice were subjected to water restriction at 1 ml per day for three days. The lever-pushing task training started 3 days after water restriction and habituation. During habituation, mice were head fixed and received water from the water tube. After starting the training, mice remained water restricted but received water during the training. Lever displacement was continuously monitored using a potentiometer, converting it into voltage signals, and recorded through a PCIe-6321 card (National Instrument). A custom LabVIEW program governed the training paradigm, precisely controlling cue presentation, reward delivery, punishment, and the determination of lever-pushing threshold crossing. Each trial was initiated with a 500 ms, 6-kHz pure tone as the cue. Mice received a water reward (approximately 8 μl) when they pushed the level surpassed the designated threshold (0.5 mm during the initial training on day 1, later increased to 1.5 mm for subsequent sessions) within the allocated task period. Failure to meet the threshold or absence of lever pushing during the task period resulted in the presentation of white noise. The inter-trial interval (ITI) was either fixed at 4 seconds or randomly varied between 3 and 6 seconds. Lever pushing during the ITI incurred an additional time-out equivalent to the ITI duration for that specific trial. The task period was 30 seconds during the first session and then reduced to 10 seconds for subsequent sessions. A total of 27 mice were trained, mice learned the task within 3 weeks, including 13 mice for calcium imaging and, 8 mice for structural imaging, and 6 mice used for behavior training.

### Movement behavior analysis

To identify movement bouts, we first determined a threshold to separate the resting and movement period. Movement bouts separated by less than 500 ms were considered continuous and were combined together^11,16^. The start time was identified as the point where the lever position crossed a threshold that exceeded the resting period, while the end time was determined by detecting the moment when the lever position fell below the threshold^11,16^. To ensure the integrity of the baseline before each movement, we adopted a specific criterion. If there were any other movements occurring within a 3-second window before a particular movement, the latter was excluded from further analysis. This exclusion step was implemented to guarantee the cleanliness and reliability of the baseline period, thus enhancing the accuracy of subsequent analyses. RM was defined as lever pushes that exceeded the threshold during the task period, while UM was those lever pushes that failed to exceed the threshold during the task period, or lever pushes during ITI.

### Activity pattern correlation and its relationship to movement trajectory correlation

Activity pattern correlation and movement trajectory correlation were calculated for each trial pair using MATLAB function ‘corrcoef’. For all trial pairs in one day, we used bins -0.2 to 0, 0 to 0.2, 0.2 to 0.4, 0.4 to 0.6, 0.6 to 0.8 and 0.8 to 1 to average all data points based on movement trajectory correlations. Then the activity pattern correlation was plotted against the movement trajectory correlation for each mouse.

### Fraction of activated ensemble difference and its relationship to movement trajectory correlation

Percentage of activated ensemble difference was calculated based on each pair of trials, if a is the number of activated bouton ensemble in trial 1, while b is the number of activated bouton ensemble in trial 2, then the fraction of activated ensemble difference for this trial pair is defined As 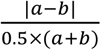, in which the |a-b| was the number of activated ensemble difference, and 0.5 ×(*a* +*b*) was the average number of activated ensemble for the trial pair. Then we calculated correlation of the movement trajectory for each trial pair using MATLAB function ‘corrcoef’. For all trial pairs in one day, we used bins -0.2 to 0, 0 to 0.2, 0.2 to 0.4, 0.4 to 0.6, 0.6 to 0.8and 0.8 to 1 to average all data points based on movement trajectory correlations. Then the percentage of activated ensemble difference was plotted against the movement trajectory correlation for each mouse.

### Image processing and analysis

For Ca^2+^ image analysis, lateral motion artifacts were corrected using the ImageJ plugin (Turboreg)^57^ or the efficient subpixel image registration algorithm^58^. Axons and boutons in FOV were manually drawn using adobe photoshop session-by-session. For the same FOV imaged both in early and late stages, only boutons with clear bouton morphology that could be identified in all sessions by visual inspection were selected and further analyzed. To extract the calcium signals for each axon or bouton, we averaged the fluorescence intensity of all labeled pixels to obtain the raw fluorescence trace. To calculate F0, we utilized a 30-second sliding window, where the 30th percentile of raw fluorescence within the window was designated as F0. ΔF/F was computed as (F-F0) / F0 for each individual axon and bouton^59^.

For structural imaging, individual boutons were identified as swellings along thinner axon shafts, and were manually identified, marked, and tracked across multiple imaging sessions using the custom written script (MATLAB). Only high-quality images displaying sparsely labeled axons, with distinct axon and bouton structures, were selected for subsequent quantification. Analysis of bouton dynamics, including formation and elimination, was performed by comparing boutons between two adjacent imaging sessions. Boutons were classified as “persistent” if they were present in both images, determined through their positions relative to nearby boutons within the same axon. An eliminated bouton was the one that appeared in the initial image but not the second image. A newly formed bouton was the one that was absent in the initial image and then appeared in the second image. The bouton survival rate was calculated as the percentage of boutons formed during day 4 of training that remained present in subsequent training sessions (days 6, 8, 10).

### Identification and classification of RM and UM axon and bouton

The activities of individual axons or boutons in both rewarded movement (RM) trials and unrewarded movement (UM) trials were aligned to the movement onset, spanning a time window from 1 second before movement initiation (served as the baseline) to 3 seconds after the movement onset. Subsequently, we calculated the average activity across all trials within this aligned time window. To identify responsive boutons, we examined the peak value of each bouton within the time window (−0.2 to 3 seconds relative to the movement onset). Boutons were considered responsive if the difference between the peak fluorescence value and the 5th percentile of the averaged activity exceeded 90% of the standard deviation (sd). For the identification of responsive axons, we plotted histograms of all peak values in RM and UM trials for each mouse. Utilizing a bin size of 0.1 sd, the peak bin values were determined for both RM and UM distributions, and the threshold was established as the mean of the corresponding peak positions in RM and UM. If the calculated threshold, based on the histogram distribution, exceeded 1 sd, the final threshold was set at 1 sd. Responsive axons were identified if the difference surpassed the threshold by comparing each axon’s peak value to the 5th percentile of the averaged activity. Subsequently, axons or boutons were categorized based on their responsiveness in RM and UM trials. Those identified as responsive exclusively in RM trials were classified as RM-only axons or boutons, while those responsive only in UM trials were categorized as UM-only axons or boutons. Axons or boutons showing responsiveness in both RM and UM trials were designated as RM-UM-both axons and boutons. To simplify, we combined the RM-only and RM-UM-both categories, grouping them as RM, RM-responsive or RM-related axons and boutons.

### Ca^2+^ event detection and identification of same or unique peaks

To detect Ca^2+^ events, we employed the Matlab findpeaks function with the following criteria: 1) z-scored ΔF/F0 exceeding 1 standard deviation^60^, and 2) raw ΔF/F0 exceeding an 8% change in fluorescence. To compare events between pairs of boutons, we considered any events occurring within 670 ms of each other as ‘matched’ and defined them as the same peak^61^, while those peaks that cannot find matched peaks were defined as unique peaks. If the same peaks or unique peaks occurred during a time window 330 ms before and 670 ms after the onset of RM or UM, those peaks were classified as RM or UM-related same or unique peaks, respectively. To calculate the same peak fraction, we divided the number of same peaks with total peaks based on each bouton pair, and averaged the results over all boutons within one axon, then averaged over all axons in one mouse.

### Principal component analysis (PCA)

We used PCA to project each trial into a lower-dimensional space to discern the low-dimensional embedding of individual boutons during rewarded movement (RM) and unrewarded movement (UM) trials. Initially, the activity of each bouton was averaged across all RM or UM trials, and the averaged activities were then concatenated for each bouton. We recorded the results in a data matrix where each column represented the concatenated trial-averaged RM and UM activity of one bouton. The size of the matrix was 2M-by-N, with M denoting the number of time points per RM or UM trial (ranging from –1 to 3 seconds relative to movement onset), and N representing the number of boutons. Subsequently, PCA was conducted across the time points of concatenated RM and UM trials, capturing the first three principal components to represent the RM and UM trials in a visually informative 3-dimensional principal component (PC) space. Each bouton was depicted as a distinct dot within this space, facilitating clear visualization and discrimination of the bouton responses during both RM and UM trials.

### PCA trajectory and calculation of selectivity index

PCA was conducted on each continuous imaged segment (4000 frames by n boutons, frame rate: 15 Hz), utilizing the first three principal components to represent the ensemble activity of boutons. Then we aligned the first three principal components from 1s before to 3 s after each RM and UM onset to generate single RM or UM neural trajectories in the PCA space. We used activity trajectory selectivity index to measure the selectivity of bouton activity towards RM or UM, a method modified from a previously published paper^62^. The activity trajectory selectivity index for an RM trial was defined as (d_to mean UM trajectory_ – d_to mean RM trajectory_) / (d_to mean RM trajectory_ + d_to mean UM trajectory_), where d_to mean UM trajectory_ (d_to mean RM trajectory_) is the Euclidean distance between the single RM trial trajectory and the mean UM (RM) trajectory, which was computed frame-by-frame. The mean RM and UM trajectories were the averages of all RM and UM trajectories respectively. For example, the first three PCs of the first frame of a RM trial is (a,b,c), while the first three PCs of the first frame of the mean UM trial is (x,y,z), then the d_to mean UM trajectory_ is 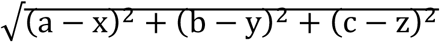. Similarly, Activity trajectory selectivity index for a UM trial was defined based on distances as (d_to mean RM trajectory_ – d_to mean UM trajectory_) / (d_to mean RM trajectory_ + d_to mean UM trajectory_). The trajectory selectivity index essentially measures how closely individual trajectories match the mean trajectories of their respective trial type versus the opposite type. For example, for an RM trial, an index score of 1 means the single trial trajectory was at the same point in PCA space as the mean RM trajectory, and an index score of -1 means the single trial trajectory was at the same point in state space as the mean UM trajectory.

### Statistics

Significance testing was performed using the Wilcoxon rank sum test, Pearson correlation coefficient, and Kolmogorov–Smirnov test using Matlab. Two-sided statistical tests were conducted, and data is presented as mean ± SEM (standard error of the mean), with all statistical tests, statistical significance values, and sample sizes described in the figure legends. Statistical thresholds used: * p < 0.05, ** p < 0.01, *** p < 0.001, NS: not significant. All source data are included in the source data table.

## Data Availability

The data, code, protocols, and key lab materials used and generated in this study are listed in a Key Resource Table alongside their persistent identifiers at Zenodo (DOI: 10.5281/ zenodo.15179158) and will be available upon publication.

## Acknowledgments

This study was supported by grants from the NINDS/NIH R01NS091144 (J.B.D.), GG gift fund (J.B.D.), Catalyst grant from The Phil & Penny Knight Initiative for Brain Resilience at the Wu Tsai Neurosciences Institute, Stanford University (J.B.D). We thank Xiaobai Ren for technical support and the members of Ding lab for valuable discussions.

## Author contributions

M.S., D.L., and J.B.D. conceptualized the project and designed the experiments. M.S., and D.L. performed animal surgeries, behavioral and in vivo imaging experiments. M.S., D.L., and K.S. analyzed the data. M.S., D.L., and J.B.D. wrote the manuscript with input from all authors.

## Reference

1. Xu, Tonghui, et al. “Rapid formation and selective stabilization of synapses for enduring motor memories.” Nature462.7275 (2009): 915–919.

2. Fu, Min, et al. “Repetitive motor learning induces coordinated formation of clustered dendritic spines in vivo.” Nature 483.7387 (2012): 92–95.

3. Yang, Guang, Feng Pan, and Wen-Biao Gan. “Stably maintained dendritic spines are associated with lifelong memories.” Nature 462.7275 (2009): 920–924.

4. Hedrick, Nathan G., et al. “Learning binds new inputs into functional synaptic clusters via spinogenesis.” Nature neuroscience 25.6 (2022): 726–737.

5. Yin, H. H., Mulcare, S. P., Hilário, M. R., Clouse, E., Holloway, T., Davis, M. I., … & Costa, R. M. (2009). Dynamic reorganization of striatal circuits during the acquisition and consolidation of a skill. Nature neuroscience, 12(3), 333–341.

6. Hwang, F. J., Roth, R. H., Wu, Y. W., Sun, Y., Kwon, D. K., Liu, Y., & Ding, J. B. (2022). Motor learning selectively strengthens cortical and striatal synapses of motor engram neurons. Neuron, 110(17), 2790–2801.

7. Graybiel, Ann M., and Scott T. Grafton. “The striatum: where skills and habits meet.” Cold Spring Harbor perspectives in biology 7.8 (2015): a021691.

8. Makino, Hiroshi, et al. “Circuit mechanisms of sensorimotor learning.” Neuron 92.4 (2016): 705–721.

9. Cataldi, Stefano, et al. “Interpreting the role of the striatum during multiple phases of motor learning.” The FEBS journal 289.8 (2022): 2263–2281.

10. Arber, Silvia, and Rui M. Costa. “Networking brainstem and basal ganglia circuits for movement.” Nature Reviews Neuroscience 23.6 (2022): 342–360.

11. Peters, A. J., Chen, S. X., & Komiyama, T. (2014). Emergence of reproducible spatiotemporal activity during motor learning. Nature, 510(7504), 263–267.

12. Oh, Seung Wook, et al. “A mesoscale connectome of the mouse brain.” Nature 508.7495 (2014): 207–214.

13. Hintiryan, Houri, et al. “The mouse cortico-striatal projectome.” Nature neuroscience 19.8 (2016): 1100–1114.

14. Hunnicutt, Barbara J., et al. “A comprehensive excitatory input map of the striatum reveals novel functional organization.” elife 5 (2016): e19103.

15. Peters, Andrew J., et al. “Striatal activity topographically reflects cortical activity.” Nature 591.7850 (2021): 420–425.

16. Sheng, M. J., Lu, D., Shen, Z. M., & Poo, M. M. (2019). Emergence of stable striatal D1R and D2R neuronal ensembles with distinct firing sequence during motor learning. Proceedings of the National Academy of Sciences, 116(22), 11038–11047.

17. Azdad, Karima, et al. “Homeostatic plasticity of striatal neurons intrinsic excitability following dopamine depletion.” PloS one 4.9 (2009): e6908.

18. Gagnon, Dave, et al. “Striatal neurons expressing D1 and D2 receptors are morphologically distinct and differently affected by dopamine denervation in mice.” Scientific reports 7.1 (2017): 41432.

19. Day, Michelle, et al. “Selective elimination of glutamatergic synapses on striatopallidal neurons in Parkinson disease models.” Nature neuroscience 9.2 (2006): 251–259.

20. De Paola, Vincenzo, et al. “Cell type-specific structural plasticity of axonal branches and boutons in the adult neocortex.” Neuron 49.6 (2006): 861–875.

21. Grillo, Federico W., et al. “Increased axonal bouton dynamics in the aging mouse cortex.” Proceedings of the National Academy of Sciences 110.16 (2013): E1514–E1523.

22. Chen, Simon X., et al. “Subtype-specific plasticity of inhibitory circuits in motor cortex during motor learning.” Nature neuroscience 18.8 (2015): 1109–1115.

23. Hasegawa, R., Ebina, T., Tanaka, Y.R., Kobayashi, K., and Matsuzaki, M. (2020). Structural dynamics and stability of corticocortical and thalamocortical axon terminals during motor learning. PLoS One 15.

24. Chen, Tsai-Wen, et al. “Ultrasensitive fluorescent proteins for imaging neuronal activity.” Nature 499.7458 (2013): 295–300.

25. Tennant, Kelly A., et al. “The organization of the forelimb representation of the C57BL/6 mouse motor cortex as defined by intracortical microstimulation and cytoarchitecture.” Cerebral cortex 21.4 (2011): 865–876.

26. Kincaid, Anthony E., Tong Zheng, and Charles J. Wilson. “Connectivity and convergence of single corticostriatal axons.” Journal of Neuroscience 18.12 (1998): 4722–4731.

27. Zheng, T., and C. J. Wilson. “Corticostriatal combinatorics: the implications of corticostriatal axonal arborizations.” Journal of neurophysiology 87.2 (2002): 1007–1017.

28. Foust, Amanda, et al. “Action potentials initiate in the axon initial segment and propagate through axon collaterals reliably in cerebellar Purkinje neurons.” Journal of Neuroscience 30.20 (2010): 6891–6902.

29. Khaliq, Zayd M., and Indira M. Raman. “Axonal propagation of simple and complex spikes in cerebellar Purkinje neurons.” Journal of Neuroscience 25.2 (2005): 454–463.

30. Clark, Beverley A., et al. “The site of action potential initiation in cerebellar Purkinje neurons.” Nature neuroscience 8.2 (2005): 137–139.

31. Doig, Natalie M., Jonathan Moss, and J. Paul Bolam. “Cortical and thalamic innervation of direct and indirect pathway medium-sized spiny neurons in mouse striatum.” Journal of Neuroscience 30.44 (2010): 14610–14618.

32. Ding, Jun, Jayms D. Peterson, and D. James Surmeier. “Corticostriatal and thalamostriatal synapses have distinctive properties.” Journal of Neuroscience 28.25 (2008): 6483–6492.

33. Albarran, Eddy, et al. “Enhancing motor learning by increasing the stability of newly formed dendritic spines in the motor cortex.” Neuron 109.20 (2021): 3298–3311.

34. Makino, Hiroshi, et al. “Transformation of cortex-wide emergent properties during motor learning.” Neuron 94.4 (2017): 880–890.

35. Luo, Liqun. Principles of neurobiology. Garland Science, 2020.

36. Hodgkin, Alan L., and Andrew F. Huxley. “A quantitative description of membrane current and its application to conduction and excitation in nerve.” The Journal of physiology 117.4 (1952): 500.

37. Debanne, Dominique, et al. “Axon physiology.” Physiological reviews 91.2 (2011): 555–602.

38. Hodgkin, Allan L., and Andrew F. Huxley. “Currents carried by sodium and potassium ions through the membrane of the giant axon of Loligo.” The Journal of physiology 116.4 (1952): 449.

39. Singer, Joshua H. “Multivesicular release and saturation of glutamatergic signalling at retinal ribbon synapses.” The Journal of physiology 580.1 (2007): 23–29.

40. Keen, Erica C., and AJ1414630 Hudspeth. “Transfer characteristics of the hair cell’s afferent synapse.” Proceedings of the National Academy of Sciences 103.14 (2006): 5537–5542.

41. Li, Geng-Lin, et al. “The unitary event underlying multiquantal EPSCs at a hair cell’s ribbon synapse.” Journal of Neuroscience 29.23 (2009): 7558–7568.

42. Chapochnikov, Nikolai M., et al. “Uniquantal release through a dynamic fusion pore is a candidate mechanism of hair cell exocytosis.” Neuron 83.6 (2014): 1389–1403.

43. Alle, Henrik, and Jorg RP Geiger. “Combined analog and action potential coding in hippocampal mossy fibers.” Science 311.5765 (2006): 1290–1293.

44. Zbili, Mickael, and Dominique Debanne. “Past and future of analog-digital modulation of synaptic transmission.” Frontiers in cellular neuroscience 13 (2019): 160.

45. Wagner, Mark J., et al. “A neural circuit state change underlying skilled movements.” Cell 184.14 (2021): 3731–3747.

46. Rueda-Orozco, Pavel E., and David Robbe. “The striatum multiplexes contextual and kinematic information to constrain motor habits execution.” Nature neuroscience 18.3 (2015): 453–460.

47. Mosberger, Alice C., et al. “Exploration biases forelimb reaching strategies.” Cell reports 43.4 (2024).

48. Masamizu, Y., Tanaka, Y.R., Tanaka, Y.H., Hira, R., Ohkubo, F., Kitamura, K., Isomura, Y., Okada, T., and Matsuzaki, M. (2014). Two distinct layer-specific dynamics of cortical ensembles during learning of a motor task. Nat. Neurosci. 17.

49. Cover, Kara K., and Brian N. Mathur. “Axo-axonic synapses: Diversity in neural circuit function.” Journal of Comparative Neurology 529.9 (2021): 2391–2401.

50. Kramer, Paul F., et al. “Synaptic-like axo-axonal transmission from striatal cholinergic interneurons onto dopaminergic fibers.” Neuron 110.18 (2022): 2949–2960.

51. Liu, Changliang, et al. “An action potential initiation mechanism in distal axons for the control of dopamine release.” Science 375.6587 (2022): 1378–1385.

52. Verdier, Dorly, James P. Lund, and Arlette Kolta. “GABAergic control of action potential propagation along axonal branches of mammalian sensory neurons.” Journal of Neuroscience 23.6 (2003): 2002–2007.

53. Kramer, Paul F., et al. “Axonal mechanisms mediating γ-aminobutyric acid receptor type A (GABA-A) inhibition of striatal dopamine release.” Elife 9 (2020): e55729.

54. Bamford, Nigel S., R. Mark Wightman, and David Sulzer. “Dopamine’s effects on corticostriatal synapses during reward-based behaviors.” Neuron 97.3 (2018): 494–510.

55. Zhai, Shenyu, et al. “Distributed dopaminergic signaling in the basal ganglia and its relationship to motor disability in Parkinson’s disease.” Current opinion in neurobiology 83 (2023): 102798.

56. Dombeck, Daniel A., et al. “Functional imaging of hippocampal place cells at cellular resolution during virtual navigation.” Nature neuroscience 13.11 (2010): 1433–1440.

57. Thevenaz, Philippe, Urs E. Ruttimann, and Michael Unser. “A pyramid approach to subpixel registration based on intensity.” IEEE transactions on image processing 7.1 (1998): 27–41.

58. Guizar-Sicairos, Manuel, Samuel T. Thurman, and James R. Fienup. “Efficient subpixel image registration algorithms.” Optics letters 33.2 (2008): 156–158.

59. Peron, Simon P., et al. “A cellular resolution map of barrel cortex activity during tactile behavior.” Neuron 86.3 (2015): 783–799.

60. d’Aquin, Simon, et al. “Compartmentalized dendritic plasticity during associative learning.” Science 376.6590 (2022): eabf7052.

61. Wagner, Mark J., et al. “Shared cortex-cerebellum dynamics in the execution and learning of a motor task.” Cell 177.3 (2019): 669–682.

62. Harvey, Christopher D., Philip Coen, and David W. Tank. “Choice-specific sequences in parietal cortex during a virtual-navigation decision task.” Nature 484.7392 (2012): 62–68.

